# Population genomic molecular epidemiological study of macrolide resistant *Streptococcus pyogenes* in Iceland, 1995-2016: Identification of a large clonal population with a *pbp2x* mutation conferring reduced *in vitro* β-lactam susceptibility

**DOI:** 10.1101/2020.05.12.090365

**Authors:** Sara B. Southon, Stephen B. Beres, Priyanka Kachroo, Matthew Ojeda Saavedra, Helga Erlendsdóttir, Gunnsteinn Haraldsson, Prasanti Yerramilli, Layne Pruitt, Luchang Zhu, James M. Musser, Karl G. Kristinsson

## Abstract

Resistance to macrolide antibiotics is a global concern in the treatment of *Streptococcus pyogenes* (Group A Streptococcus, GAS) infections. In Iceland, since the detection of the first macrolide-resistant isolate in 1998, three epidemic waves of macrolide-resistant GAS infections have occurred with peaks in 1999, 2004, and 2008. We conducted whole genome sequencing of all 1,575 available GAS macrolide-resistant clinical isolates of all infection types collected at the national reference laboratory in Reykjavik from 1998 to 2016. Among 1,515 erythromycin-resistant isolates, 90.3% were of only three *emm* types: *emm4* (*n* = 713), *emm6* (*n* = 324), and *emm12* (*n* = 332), with each being predominant in a distinct epidemic peak. The antibiotic efflux pump genes, *mef*(A) and *msr*(D), were present on chimeric mobile genetic elements in 99.3% of the macrolide-resistant isolates of these *emm* types. Of note, in addition to macrolide resistance, virtually all *emm12* isolates had a single amino acid substitution in penicillin-binding protein PBP2X that conferred a two-fold increased penicillin G and ampicillin MIC among isolates tested. We conclude that each of the three large epidemic peaks of macrolide-resistant GAS infections occurring in Iceland since 1998 was caused by the emergence and clonal expansion of progenitor strains, with macrolide resistance being conferred predominantly by inducible Mef(A)-Msr(D) drug efflux pumps. The occurrence of *emm12* strains with macrolide resistance and decreased beta-lactam susceptibility was unexpected and is of public health concern.

**Repositories:** Genomic sequencing data for all 1,515 macrolide/erythromycin-resistant isolates were deposited into the National Center for Biotechnology Information Sequence Read Archive under bioproject accession PRJNA614628 and assembled sequences for composite elements Φ29854, Φ29862 and Φ29661 were deposited in Genbank under accessions ###, ### and ### respectively.

## INTRODUCTION

*Streptococcus pyogenes* (group A streptococcus, GAS) is an important human pathogen that globally is among the top 10 infectious causes of human mortality, causing over 700 million infections and almost 520,000 deaths annually (1, 2). GAS causes a wide spectrum of diseases ranging from prevalent uncomplicated mild infections such as pyoderma (111 million cases/yr) and pharyngitis (616 million cases/yr), to relatively infrequent severe life-threatening invasive infections such as necrotizing fasciitis/myositis and sepsis (663,000 cases/yr causing 163,000 deaths/yr) (1, 2). GAS produces a myriad of extracellular virulence factors that contribute to adhesion, degradation and breaching of tissue barriers, subversion and evasion of host innate and adaptive immune defenses, and systemic intoxication, among many other pathogenic processes (3–5). Among these the Emm/M protein encoded by the *emm* gene is a major virulence factor with multiple functions, including promoting adherence to human epithelial cells and inhibiting phagocytosis in the absence of opsonizing antibodies (6). The M protein is the primary surface antigen eliciting the human immune response. Diversification in the first 180 nucleotides of the *emm* gene encoding the hypervariable amino-terminus of the M protein, is the basis for *emm* typing, the most commonly used epidemiological marker of *S. pyogenes* strain lineages (7). There are over 250 *S. pyogenes emm* types listed in the CDC *emm* database as of September 11, 2019 (8). Importantly, there is no licensed vaccine to prevent *S. pyogenes* infections (9).

Beta-lactam antibiotics that inhibit peptidoglycan synthesis are the primary antibacterial treatment for *S. pyogenes* infections, and despite over 75 years of use, no penicillin-resistant clinical isolate has been reported (10, 11). However, two recent studies reporting reduced *in vitro* susceptibility to beta-lactam antibiotics among clinical isolates are of concern (12, 13). Macrolides are secondary alternative antibiotics recommended for individuals allergic to penicillin. Macrolides, and the mechanistically similar lincosamides and streptogramins, inhibit protein translation through binding interactions with the ribosome. Because of drug synergism and the potential benefits of inhibiting extracellular protein/toxin production, combination antibiotic therapy of a beta-lactam and a lincosamide (e.g. penicillin and clindamycin) is recommended for severe invasive *S. pyogenes* infections. In *S. pyogenes* there are two principal mechanisms for acquired macrolide resistance: target site modification and active efflux (14, 15). Target site modification is mediated by <u>e</u>rythromycin r<u>R</u>NA <u>m</u>ethylases, predominantly Erm(B) and Erm(TR), which methylate the 23S rRNA and block antibiotic binding to the ribosome. This modification provides resistance to <u>m</u>acrolides, <u>l</u>incosamides, and <u>s</u>treptogramin <u>B</u> and confers the MLS_B_ resistance phenotype. Active efflux is mediated by proton-dependent membrane-associated pumps that transport 14- and 15-membered macrolides out of the bacterial cell (but not 16-membered macrolides, lincosamides, or streptogramins), conferring the M resistance phenotype. Although the <u>m</u>acrolide <u>ef</u>flux activity was initially attributed to <u>Mef</u>(A (16), recent *mef*(A) and *msr*(D) gene knock-out and knock-in experiments demonstrate that Msr(D) is the functionally predominant macrolide efflux transporter in *S. pyogenes* strains of multiple *emm* types (17, 18). Macrolide resistance genes are not part of the GAS core chromosome but are acquired, encoded largely on a diverse set of integrative conjugative elements and chimeric mobile genetic elements (MGE), such as those formed by the integration of an ARG-encoding transposon into a prophage (19–22). Resistance to macrolides at low frequency can also spontaneously arise via mutations in the 23S rRNA and in ribosomal proteins L4 and L22, encoded by genes *rplD* and *rplV*, respectively (14).

Since the first reports of macrolide-resistant GAS in England in the late 1950s (23), resistance has disseminated worldwide, and its prevalence has been reported to vary profoundly geographically (i.e. between countries/regions at a point in time) and temporally (i.e. in the same country/region over time) (15, 24). In many instances, an increase in the prevalence of resistant isolates clearly corresponded with increased antibiotic usage, consistent with the influence of antibiotic selective pressure (25). However, in some cases precipitous changes in resistance prevalence has occurred in association with a change in predominant GAS clone or mechanism of resistance, but independent of any perceived change in antibiotic usage (26). In Iceland, erythromycin susceptibility testing was performed on at least 100 GAS isolates per year and the first macrolide-resistant isolate was not detected until early 1998. Over the next year the monthly proportion of macrolide-resistant GAS precipitously increased from 0% in March 1998 to 56% in March 1999 (27). Among 367 erythromycin-resistant GAS isolates collected through July 1999, 99% were M resistance phenotype. T-antigen typing of 30 isolates collected from July to December 1998 revealed 3 T-types: T8 (73%), T6(17%), and T28(10%). Among 67 isolates compared by SfiI restriction pulse-field gel electrophoresis (PFGE), 58 had the same banding pattern (27). The finding that the majority of the isolates were T8 and of a single PFGE pattern suggested the 1999 epidemic wave was likely mono- or pauci-clonal in nature. Of note, over the same time frame (c.a. 1998-2001) a significant increase in macrolide-resistant GAS also occurred in Spain (28) and in Toronto, Canada (29). A second modest peak of increased macrolide-resistant GAS in Iceland occurred in 2004, followed by a third, larger and rapidly-arising peak in 2008 (30). Here we present whole genome sequencing-based molecular epidemiological characterization of all available *S. pyogenes* erythromycin-resistant isolates (*n* = 1,515) collected in Iceland from 1995 to 2016. Emphasis is placed on the three predominant *emm* types (4, 6, and 12) causing the three successive epidemic peaks of macrolide-resistant infections.

## METHODS

### Bacterial Isolates

A total of 15,217 GAS strains were isolated from patient specimens submitted to the Department of Clinical Microbiology, Landspitali University Hospital, from 1995 to 2016. The laboratory receives invasive (e.g. blood and CSF) isolates from the whole country and acts as the primary laboratory for other GAS cultures for about 75% of the country. Macrolide-resistant isolates were stored in glycerol broths at −85°C (invasive isolates) or −20°C (non-invasive isolates). The majority of the samples (~60%) were collected from patients from the Reykjavík capital region. According to Iceland Statistics (hagstofa.is.en), the population of Iceland and Reykjavík in 1995 was 267,809 and 158,597, respectively. In 2016, the population of Iceland was 332,529 and in the Reykjavík region was 209,500. Information regarding the 15,217 isolates (e.g. sample origin, geographic place of collection, date of collection, antibiotic susceptibility, patient residence, and patient age and gender) was recorded in the Laboratory Information System at the Department of Clinical Microbiology. Since 1998, all GAS detected in the department have been collected and stored frozen. Isolates, except those from urine samples, were tested for erythromycin susceptibility by the disk diffusion method based on CLSI criteria (31), and after June 2012 based on methods and criteria from EUCAST (32). Isolates were considered to be the same strain if they were collected twice or more, ≤7 days apart from the same patient. When antibiotic resistant susceptibilities were inconsistent between isolates taken from the same patient, the isolate from the more invasive infection sample was used. Isolates were grown on tryptic soy agar with 5% sheep blood (Benton Dickson) or with 5% horse blood (Oxoid) at 37°C and 5% CO_2_.

### Whole Genome Sequencing

All viable GAS isolates that tested resistant to the macrolide antibiotic erythromycin (*n* = 1,575) within the collection were sent to the Center for Molecular and Translational Human Infectious Diseases Research, Department of Pathology and Genomic Medicine, Houston Methodist Research Institute (Houston, Texas) for whole genome sequencing. Genomic DNA extraction and multiplexed library preparation was performed as previously described (33). Paired-end, 150 nucleotide-long sequencing reads were obtained using an Illumina NextSeq 500 sequencer. Sequence data preprocessing (i.e. artifact and adapter trimming, quality filtering, and base call error correction) and *de novo* assembly for each isolate was done as previously described (33).

### Initial genetic typing and gene content profiling

Multi-locus sequence type (MLST), *emm* type, and antibiotic resistance gene content was determined for each isolate from the sequencing reads relative to publicly available reference databases using SRST2 software (34) as previously described (33). Mobile genetic element typing was determined relative to a published database of *S. pyogenes* phage and ICE-encoded integrase and virulence factor genes as previously described using SRST2 (35). The *pbp2x* gene was identified in and retrieved from isolate genome assemblies using blastn and bedtools-getfasta respectively.

### Polymorphism discovery

Sequence reads were mapped to relevant reference sequences using SMALT (www.sanger.ac.uk/resources/software/smalt), and polymorphisms between the aligned reads and the reference sequences were identified using FreeBayes (36). Polymorphisms were filtered on the basis of call consensus (≥70%), mapped quality (≥Q30), and coverage depth (≥10-fold) using VCFlib (www.github.com/ekg/vcflib#vcflib). Specifically, for *emm4* isolates, core chromosomal single nucleotide polymorphisms (SNPs) were called relative to the genome of strain MGAS10750 (20), *emm6* isolates to MGAS10394 (19), and *emm12* to MGAS9429 (20). SNPs were annotated, and the effects of variants were predicted using SnpEff (37). Polymorphisms in the chimeric elements encoding *mef*(A)-*msr*(D) among *emm4, emm6*, and *emm12* strains were called relative to Φ29862, Φ29961, and Φ29854 respectively.

### Phylogenetic inference and population structure

Concatenated SNP sequences used for evaluation of genetic relationships among isolates were generated using Prephix and Phrecon (www.github.com/codinghedgehog). To limit phylogenetic inferences to primarily vertically inherited core chromosomal SNPs, mobile genetic element (phage and ICE) encoded regions were excluded and regions of horizontal transfer and recombination were identified and excluded using Gubbins (38). Phylogeny among isolates was inferred by the Neighbor-Joining method using SplitsTree (39), and phylograms were generated with Dendroscope (40). Genetic distances among the isolates were calculated using MEGA 7 (41).

### Construction of isogenic strain with PBP2X-Met593Thr variant

Strain MGAS27213-L601P, M593T was constructed from MGAS27213-L601P by allelic exchange using methods previously described with modifications (42). Overlap extension PCR was used to introduce the Met593Thr substitution into *pbp2x* of MGAS27213-L601P. Primers *pbp2x*-5’fwd (CAATTGTACAAAACCGTTACGATCC AAG) and *pbp2x*-5’rev (TAGTAACATACATCAAAAAGTCTGGTTTATC) were used to amplify the *pbp2x* 5’ end. Primers *pbp2x*-T593-3’fwd (CTTTTTGATGTATGTTACTACGACTAAACCAC) and *pbp2x-* 3’rev (GTGAATACATATCAGTATTTGTGGGTCATC) were used to amplify the *pbp2x* 3’end introducing a single A to C nucleotide change in *pbp2x* condon 593. Primers pBBL740-fwd (GTAACGGTTTTG TACAATTGCTAGCGTAC) and pBBL740-rev (AAATACTGATATGTATTCACGAACGAAAATC) were used to amplify and linearize suicide plasmid pBBL740 by inside-out PCR. The *pbp2x* 5’-end and 3’-end amplicons were spliced with the linearized pBBL740 amplicon using NEBuilder HiFi kit (New England Biolabs). The resultant spliced plasmid was transformed into parental strain MGAS27213-L601P and single cross-over transformants were selected by plating on THY agar with chloramphenicol 10ug/ml. Transformants were screened by genomic DNA PCR amplification and Sanger sequencing using primers *pbp2x*-5’fwd and *pbp2x*-seq (GATGTCTCACCAG GATTCTTTC). Ten confirmed single cross-over transformants were pooled, expanded by outgrowth and then passaged 8 times on THY agar plates without chloramphenicol to promote double cross-over resolution. Chloramphenicol sensitive isolates were identified by duplicate plating and screened for the *pbp2x*-Thr593 allelic exchange by PCR amplification and Sanger sequencing. Resultant candidate MGAS27213-L601P, M593T derived strains were whole genome sequenced to confirm the lack of spontaneous spurious mutations.

## RESULTS

### Epidemiological surveillance

In Iceland, 15,217 beta-hemolytic group A carbohydrate antigen-positive streptococcal clinical isolates were detected from patients with noninvasive and invasive infections from 1995 to 2016 (Figure 1). The majority of the isolates, 10,010 (66%), were from the upper respiratory tract, nearly all (95%) of which were from throat. Of the 15,217 isolates, 1,806 (11,9%) were macrolide resistant, of which 1,515 (83.9%) were stored and viable upon retrieval. Isolation sites included: upper respiratory tract, 1,137 (75.0%); skin/wound, 214 (14.1%); middle ear, 92 (6.1%); lower respiratory tract, 24 (1.6%); abscess, 21 (1.4%); blood, 11 (0.7%); and other, 16 (1.1%). The proportion of available macrolide-resistant isolates per year ranged from 35.2% for 2003 to 98.0% for 1999 (Figure 1).

**Figure 1.**
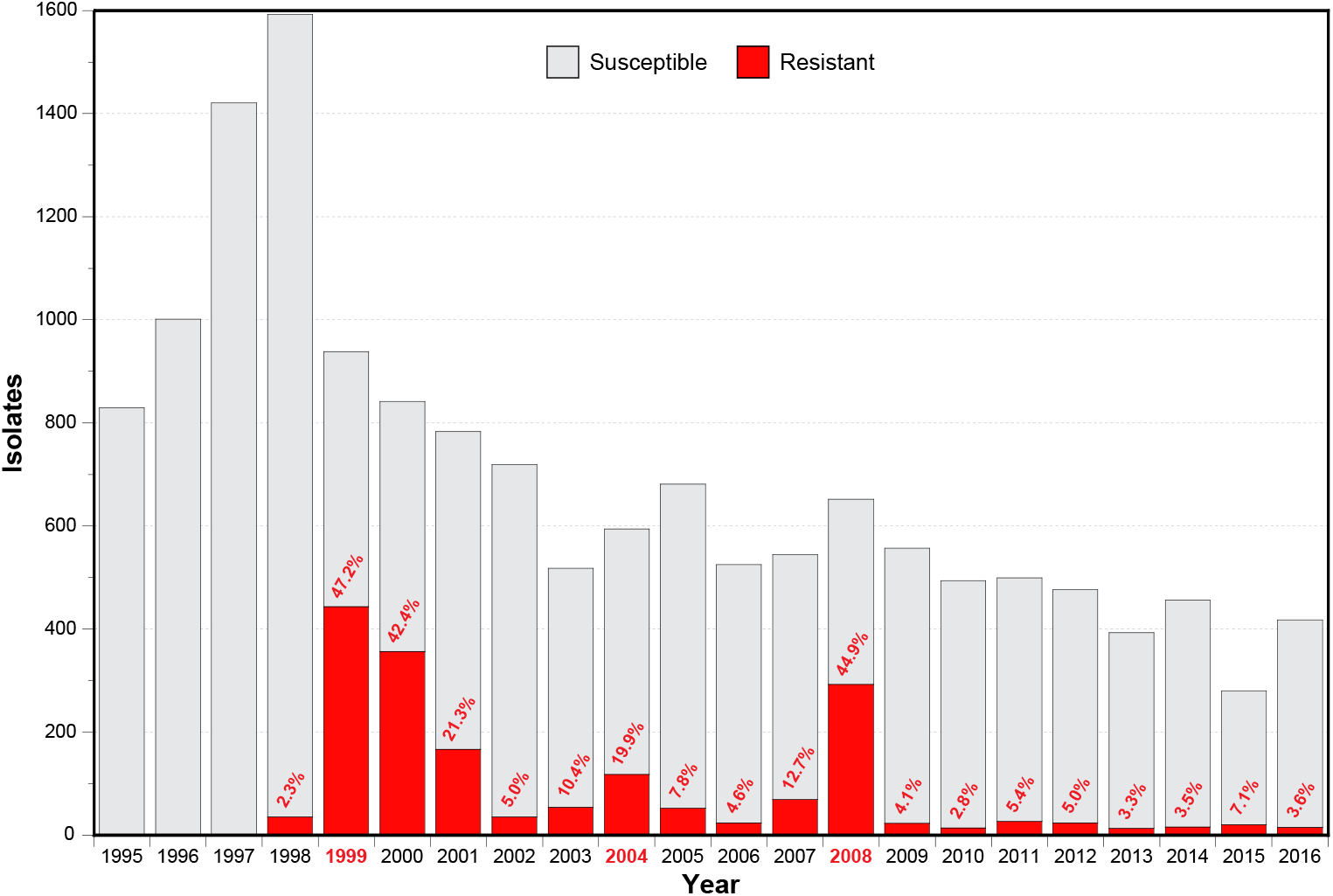
Annual incidence of positive GAS cultures at the Landspitali University Hospital from 1995 to 2016. The proportion of erythromycin susceptible and resistant isolates are colored as indicated in the index. The annual percent of erythromycin-resistant isolates is provided.

No macrolide-resistant GAS isolate was identified until July 1998. Following this, resistant isolates rapidly increased in proportion to a peak of 47.2% of isolates in 1999. Resistant isolates gradually declined in proportion to 5.0% in 2002. A second more modest increase in the proportion of resistant isolates peaked at 19.9% in 2004. A third peak of resistance rose to 44.9% of GAS isolates in 2008. In total, 1,802 of 15,217 (11.2%) of GAS clinical isolates tested were resistant to erythromycin (Figure 1).

### Whole genome sequencing genetic characterization

To genetically characterize the cohort, all 1,575 available viable erythromycin-resistant GAS isolates were whole genome sequenced to an average 214-fold depth of coverage (range: 18 to 1859×) using Illumina paired-end sequencing. Based on the sequence data, 60 of the isolates were excluded from the investigation, for reasons such as the isolate was not *S. pyogenes*, was a duplicate, or was contaminated. The retained 1,515 erythromycin-resistant *S. pyogenes* isolates and their epidemiological and genetic characteristics are listed in Table S1. Sequence reads for the isolates assembled on average into 67 contigs summing to 1.82 Mbp with a G+C content of 38.4%, values which are consistent with closed genomes of *S. pyogenes*.

The 1,515 macrolide-resistant isolates were comprised of 27 *emm* types (Table S1 and Figure 2). Three *emm* types, *emm4* (*n* = 713, 47.1%), *emm12* (*n* = 332, 21.9%), and *emm6* (*n* = 324, 21.4%), account for the majority of the isolates (*n* = 1,369, 90.4%). Analysis of the epidemic curve by *emm* type shows the first wave (years 1998-2001) of macrolide-resistant isolates was composed predominantly of *emm4* (74%) with some *emm12* (24%). The second wave (years 2004-2005) was composed predominantly of *emm12* (68%) with some *emm75* (17%). And the third wave (years 2007-2008) was composed predominantly of *emm6* (91%).

**Figure 2.**
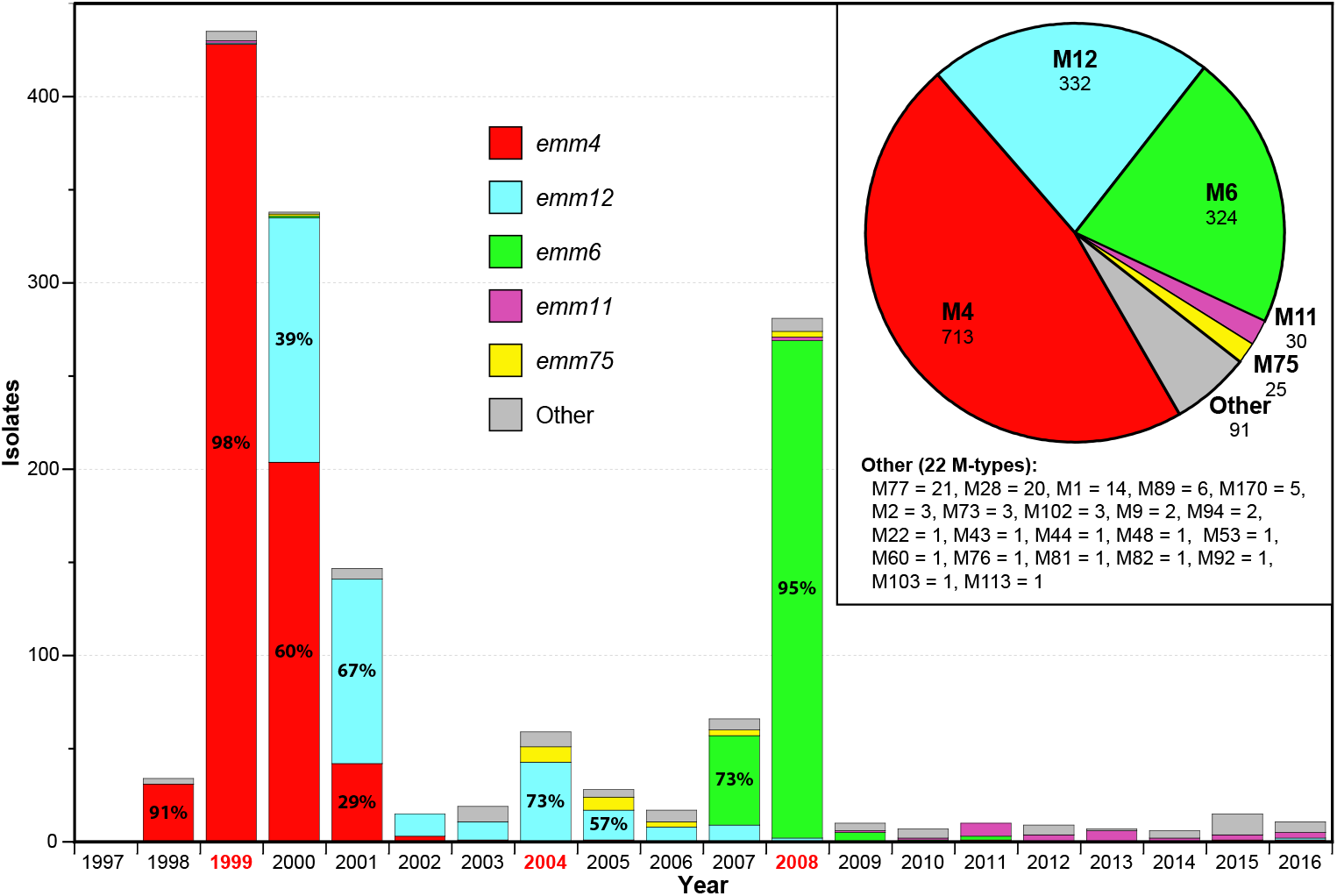
Annual incidence of erythromycin-resistant isolates with the number and proportion of *emm* types. Isolate proportions are colored by *emm* type as indicated in the index. The inset pie chart illustrates the 1,515 erythromycin-resistant isolates proportioned by *emm* type. The five most prevalent *emm* types shown (4, 12, 6, 11, and 75 in decreasing order) account for 94.% of the isolates.

Analysis of the antibiotic resistance gene (ARG) content of the cohort identified 17 different ARGs that were present in 21 different combinations (Table 1). One or more ARG was detected in 1,471 (97.1%) of the isolates, and no macrolide-resistant gene was found in 44 isolates of 13 different *emm* types (Table S1). Previous publications have shown that *emm* types 4, 6, 12, and 75 are commonly associated with macrolide resistance. The most prevalent combination of macrolide resistance genes was *mef*(A)-*msr*(D), conferring the M resistance phenotype, was found in 1,369 (90.4%) isolates. Virtually all isolates (1,359/1,369, 99.3%) of the three most prevalent *emm* types (4, 6, and 12) encode *mef*(A)-*msr*(D) (Figure 3). Reciprocally, virtually all (1,359/1,369, 99.3%) isolates encoding *mef*(A)-*msr*(D) in the cohort are of *emm* types 4, 6, or 12. The *erm*(B) gene was found in 69 (4.6%) isolates (43/69 = *emm* types 11 and 75) and the *erm*(TR) gene in 30 (2.0%) isolates (20/30 = *emm77*). Thus, 99 (6.5%) of the isolates have an erythromycin rRNA methylase gene conferring the MLS_B_ phenotype. No isolate was found that encoded both a macrolide efflux and an erythromycin resistance methylase gene.

**Figure 3.**
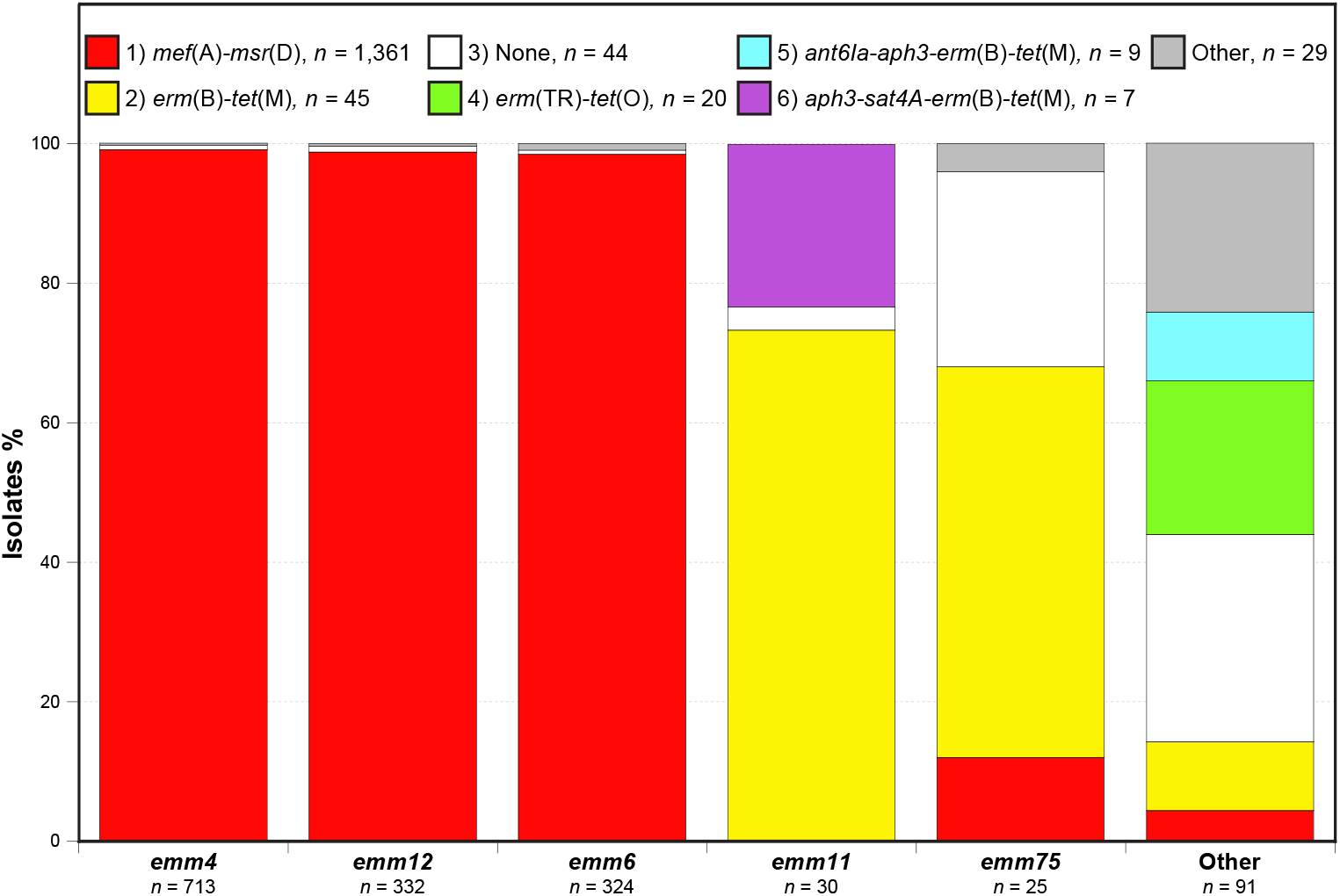
Antibiotic resistance genes and their proportions according to *emm* type, colored as indicated in the index. The six most prevalent ARG content combinations account for 98.5% of the 1,515 erythromycin-resistant isolates.

**Table 1.**
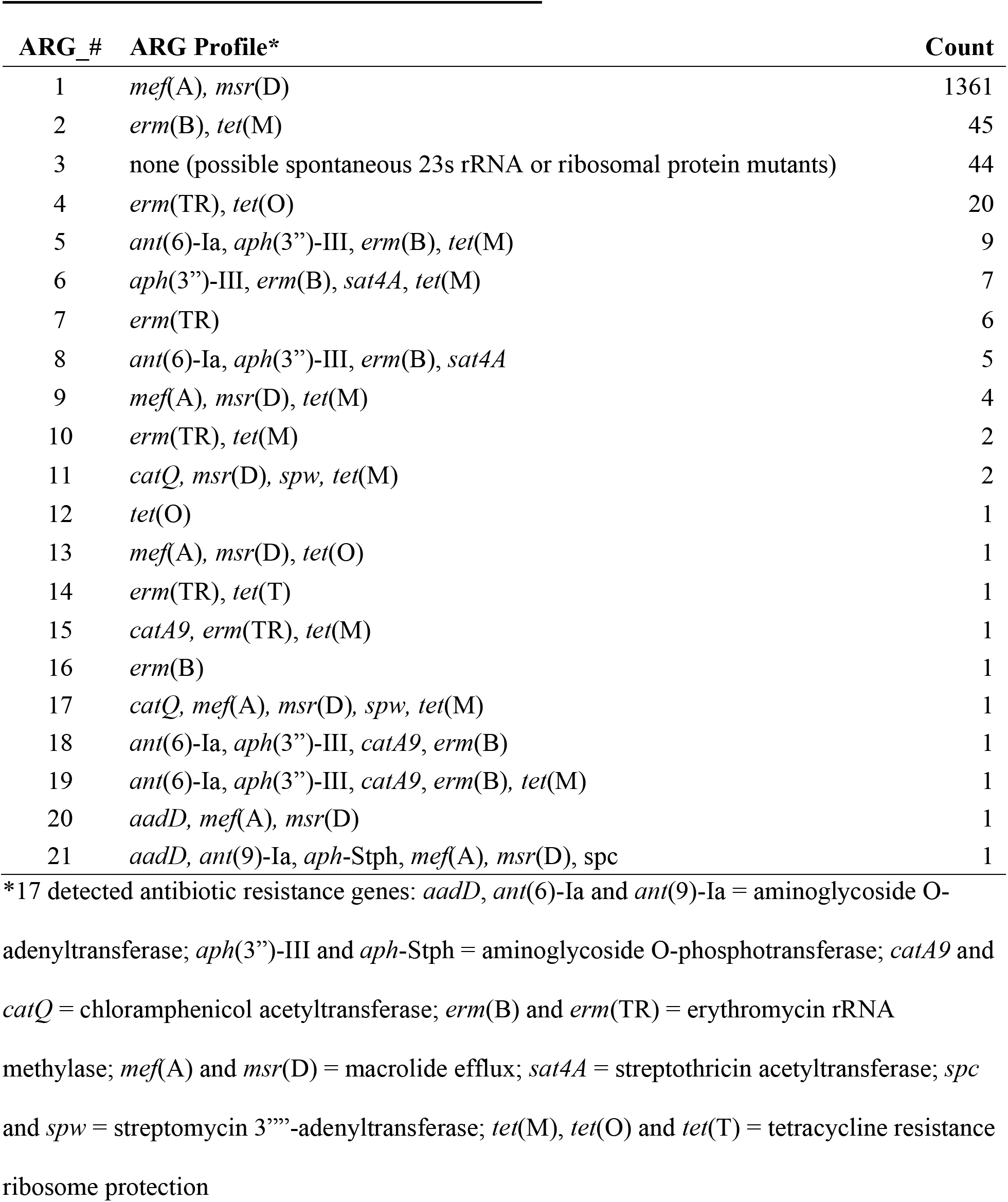
Antibiotic Resistance Genes and Profiles.

### Phylogenetic relationships

Within an *emm* type, the macrolide-resistant *mef*(A)-*msr*(D)-encoding isolates are closely genetically related, consistent with the isolates arising from clonal expansion of a recent common progenitor (Figure 4). The *mef*(A)-*msr*(D)-encoding *emm4* (*n* = 709), *emm6* (*n* = 322), and *emm12* (*n* = 327) isolates across the 1.7 Mbp core chromosome differed pairwise on average by only 11.3, 9.2, and 20.8 SNPs, respectively. For each of these *emm* types, the few erythromycin resistant isolates that lacked any detectable ARGs were more genetically distant from the *mef*(A)-*msr*(D)-encoding isolates and appear to represent infrequent sporadic spontaneous resistant mutants.

**Figure 4.**
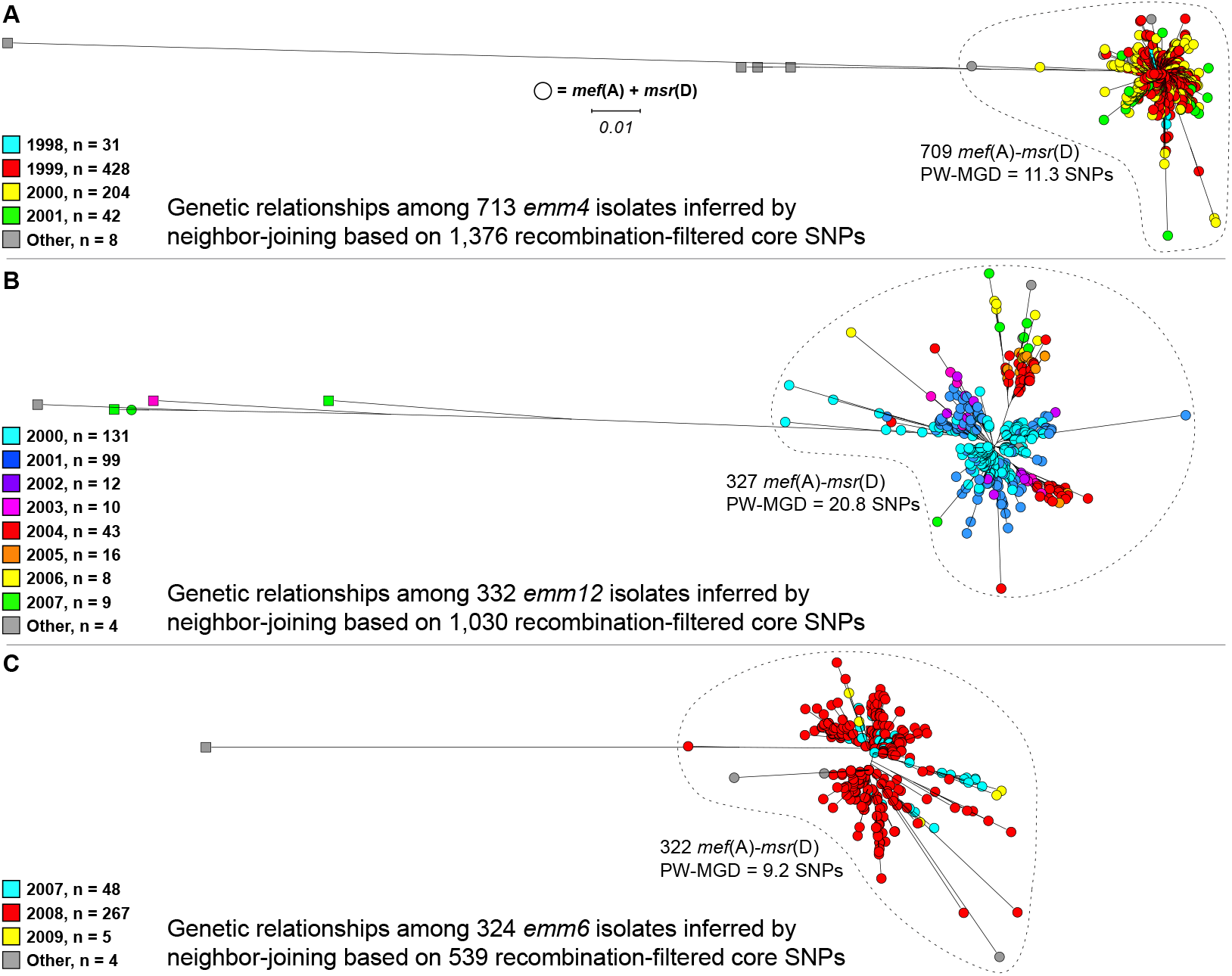
Genetic relationships among erythromycin-resistant isolates. Illustrated at the same scale are trees for the three most prevalent *emm* types that account for 90.3% of the 1,515 detected erythromycin-resistant isolates. Isolates encoding *mef*(A) and *msr*(D) are shown with circles and isolates that do not are shown with squares. Closely-related clonal isolates encoding *mef*(A) and *msr*(D) are enclosed within dotted lines. The isolates are colored by year of detection as indicated. A) Phylogeny inferred for *emm4* isolates. B) Phylogeny inferred for *emm12* isolates. C) Phylogeny inferred for *emm6* isolates.

To investigate the context of the *mef*(A)-*msr*(D) genes within the isolate genomes, the *de novo*-assembled contigs of the *emm* type 4, 6, and 12 isolates were searched using BLASTn. For each of these *emm* types, the *mef*(A) and *msr*(D) genes were found adjacently encoded on transposon Tn1207.1 inserted into a phage forming a composite MGE like that first described for Φ10394.4 of macrolide-resistant *emm6* strain MGAS10394 (19, 43). These elements were all found integrated at the same site in the genome disrupting the *comEC* gene. Full length *de novo* assemblies of the *mef*(A)-*msr*(D)-encoding MGEs were obtained from *emm4* strain MGAS29862 (Φ29862 = 52479 bp, 57 CDSs), *emm6* strain 29961 (Φ29961 = 57866 bp, 57 CDSs), and *emm12* strain MGAS29854 (Φ29854 = 52542 bp, 58 CDSs). The sequences of these elements share high identity (>98%) with each other and to Φ10394.4 (Figure S1).

Accurately detecting SNPs in phages in the *S. pyogenes* genome is problematic as most isolates are polylysogenic, which frequently causes erroneous cross-mapping of reads and to overcalling SNPs in phage. Despite this, mapping the whole genome sequencing reads of the erythromycin-resistant isolates to the *mef*(A)-*msr*(D)-encoding MGEs detected relatively few SNPs. The 709 *emm4* isolates differed pairwise by 0.4 SNPs determined relative to Φ29862, the 322 *emm6* isolates by 4.6 SNPs relative to Φ29961, and 326/328 (99%) *emm12* isolates by 0.95 SNPs relative to Φ29854. The finding that the isolates of the same *emm* type have *mef*(A)-*msr*(D)-encoding composite MGEs that are nearly sequence invariant is again consistent with the macrolide-resistant isolates stemming from recent clonal expansions.

Comparison of *S. pyogenes* genomes has identified strain-to-strain differences in MGE content stemming from the dynamic gain and loss of ICEs and phages as the largest source of genetic diversity. As a third measure of relatedness, the MGE content of the isolates was assessed by sequence read mapping relative to a database of known *S. pyogenes* MGE-encoded integrases (*n* = 31) and virulence factors (*n* = 19). This comparison process generates a 50-allele present/absent genotype. Among the 322 erythromycin-resistant *emm6* isolates, 309 (96%) have the same inferred MGE content (Table S2), as do 309 of the 327 (95%) *emm12* isolates. Among the 709 erythromycin-resistant *emm4* isolates, 690 (97%) have the same inferred MGE-encoded virulence factor content, although they differed more extensively in the detected MGE integrase gene content. Results of the analysis of MGE-encoded gene content was consistent with the SNP data for the core chromosome and the *mef*(A)-*msr*(D)-encoding composite MGEs. Our data demonstrate the erythromycin-resistant *emm* type 4, 6, and 12 isolates are within their respective *emm* type, each closely genetically related, consistent with the isolates of each *emm* type stemming from recent clonal expansions.

### Potential for altered beta-lactam antibiotic susceptibility

Recently it has been shown that many *S. pyogenes* clinical isolates with nonsynonymous (amino acid substituting) nucleotide changes in the penicillin-binding protein 2X gene (*pbp2x*) are associated with reduced susceptibility *in vitro* to one or more members of the beta-lactam family of antibiotics (12, 13). Among the 1,515 macrolide-resistant isolates, 25 *pbp2x* alleles encoding for 10 PBP2X variants were identified (Table S3). Although the *pbp2x* allele differed from one *emm* type to another, virtually no allelic variation in *pbp2x* was found within an *emm* type for the cohort. That is, in terms of *pbp2x* allele/PBP2X variants, 712/713 *emm4* isolates have the same *pbp2x* allele/PBP2X variant, 324/324 *emm6* are the same, and 327/332 *emm12* are the same. The *emm4* isolates have the consensus PBP2X wild-type (WT) sequence that is most prevalent among *S. pyogenes* isolates of multiple *emm* types (12). The PBP2X variant of all 324 *emm6* isolates have three substitutions (Ile502Val, Pro676Ser, and Lys708Glu) and 327/332 *emm12* isolates have a single substitution (Met593Thr) relative to the PBP2X WT 751 amino acid sequence. This lack of *pbp2x* sequence diversity is again consistent with *emm* type 4, 6, and 12 macrolide-resistant isolates stemming from recent clonal expansions.

The susceptibility to penicillin G, ampicillin, and erythromycin of the three predominant PBP2X variants present in the *emm* 4, 6, and 12 isolates was tested for five isolates of each *emm* type (Table 2). The isolates were selected to represent the temporal spread of each *emm* type corresponding with the three peaks of macrolide-resistant infections. All five *emm4* isolates having the PBP2X WT variant were fully susceptible to the beta-lactam antibiotics penicillin G and ampicillin. Despite the five *emm6* isolates having a PBP2X variant with three amino acid substitutions relative to the PBP2X WT, they were also fully susceptible to the beta-lactam antibiotics. In contrast, all five *emm12* isolates with a Met593Thr substitution PBP2X variant had approximately two-fold increased MICs for both penicillin G and ampicillin. To unambiguously determine if the PBP2X Met593Thr substitution alters beta-lactam susceptibility, we constructed an isogenic PBP2X Thr593 substitution derivative using the parental strain MGAS27213-PBP2X-L_601_P (12). Importantly, whole genome sequencing confirmed that the constructed derivative strain, MGAS27213-PBP2X-L_601_P,M_593_T, only differs from the parent strain by a single nucleotide change in codon 593 (<u>A</u>TG to <u>C</u>TG) of *pbp2x*. As anticipated, the parental strain had fully susceptible PBP2X WT penicillin G and ampicillin MIC levels. In contrast, the isogenic PBP2X Met593Thr derivative had two-fold increased MICs (Table 2). All 15 of the *emm4, emm6*, and *emm12* isolates encoding *mef*(A)-*msr*(D) were erythromycin resistant, and both the parental and PBP2X Met593Thr derivative strains were erythromycin susceptible.

**Table 2.** Antibiotic Minimal Inhibitory Concentrations.

| *emm* | Isolate | Date | Peak | PBP2X Substitutions* | Penicillin-G<br>0.002-32 ug/ml<sup>†</sup> | Ampicillin<br>0.016-256 ug/ml<sup>†</sup> | Erythromycin<br>0.016-256 ug/ml<sup>†</sup> |
| --- | --- | --- | --- | --- | --- | --- | --- |
| 4 | MGAS31145 | Feb-1998 | 1 | Consensus WT | 0.012 | 0.016 | 12 |
|  | MGAS30167 | Jan-1999 | 1 | Consensus WT | 0.012 | 0.016 | 12 |
|  | MGAS31312 | Jun-1999 | 1 | Consensus WT | 0.012 | 0.016 | 12 |
|  | MGAS30569 | Jan-2000 | 1 | Consensus WT | 0.016 | 0.016 | 8 |
|  | MGAS29862* | Oct-2001 | 1 | Consensus WT | 0.012 | 0.016 | 12 |
| 12 | MGAS30669 | Jun-2000 | 1 | M593T | 0.023 | 0.032 | 12 |
|  | MGAS29854* | Apr-2001 | 1 | M593T | 0.023 | 0.023 | 12 |
|  | MGAS29776 | May-2003 | 2 | M593T | 0.023 | 0.032 | 12 |
|  | MGAS31113 | Jun-2004 | 2 | M593T | 0.023 | 0.032 | 8 |
|  | MGAS31135 | Jul-2005 | 2 | M593T | 0.023 | 0.032 | 12 |
| 6 | MGAS30249 | Aug-2007 | 3 | I502V, P676S, K708E | 0.012 | 0.016 | 8 |
|  | MGAS30277 | Jan-2008 | 3 | I502V, P676S, K708E | 0.016 | 0.016 | 8 |
|  | MGAS29961* | Apr-2008 | 3 | I502V, P676S, K708E | 0.012 | 0.016 | 8 |
|  | MGAS30512 | Nov-2008 | 3 | I502V, P676S, K708E | 0.012 | 0.016 | 8 |
|  | MGAS30516 | Feb-2010 | 3 | I502V, P676S, K708E | 0.016 | 0.016 | 8 |
| 89§ | MGAS27213-PBP2X-L601P | - | - | S562T | 0.016 | 0.016 | 0.125 |
|  | MGAS27213-PBP2X-L601P,M593T | - | - | S562T, M593T | 0.032 | 0.032 | 0.125 |

\*Amino acid substitutions relative to PBP2X consensus WT sequence
<sup>†</sup>Antibiotic concentration range of gradient method E-test strips.
‡ Isolates from which composite MGEs Φ29862, Φ29854 and Φ29661 encoding *mef(A)-msr(D)* were assembled.
§ Reference strains used for construction of isogenic *pbp2x* alleles/PBP2X variants, not part of the Iceland macrolide resistant cohort

## DISCUSSION

Macrolide-resistant GAS first appeared in Iceland in 1998 and has for most years since been relatively rare, with a yearly incidence typically below 5%. This contrasts with three rapid increases reaching peaks in 1999 (47.2%), 2004 (19.9%), and 2008 (44.9%). These peaks suggested clonal epidemics, now confirmed in this study. The first wave (1998-2001) was composed predominantly of *emm4* (74%), the second (2004-2005) of *emm12* (68%), and the third (2007-2008) of *emm6* (91%). The peaks neither coincided with significant changes in either the type or amount of macrolide consumed over the year preceding the peaks. This suggests that GAS clones can spread rapidly in populations where herd immunity may be low to that particular clone, decline in numbers as herd immunity increases, and be replaced by another newly-emerging clone. The results presented here should not be interpreted as macrolide consumption having had no effect on the three epidemic peaks, but alternatively that significant changes in macrolide usage are not necessary for there to be significant changes in the prevalence of macrolide-resistant GAS infections and the predominant clone causing such infections. Similar results showing a lack of correspondence between macrolide consumption and occurrence of macrolide-resistant GAS isolates in Portugal has been reported, where decline in erythromycin resistance was associated with the disappearance of isolates belonging to an *emm3*-ST315 lineage and yet accompanied by a high consumption of macrolides (26).

The short time between the emergence in Iceland of the first erythromycin-resistant *emm4* isolates in 1998 and the first *emm12* isolates in 2000 (Figure 1), with both contributing to the first macrolide-resistant epidemic wave (1998-to-2001), and the similarity of the *mef*(A)-*msr*(D) encoding elements in these *emm* types (Figure S1), raises the possibility that the emergence events are directly related. That is, it is possible that the *emm12* lineage progenitor arose through recent horizontal acquisition of the *mef*(A)-*msr*(D) composite MGE directly from an Icelandic *emm4* donor. Alignment of *emm4* Φ29862 with *emm12* Φ29854 revealed a difference of 668 SNPs. The several hundred-fold greater numbers of SNPs identified for the *mef*(A)-*msr*(D) composite MGEs inter *emm* type vs intra (<1 SNP pairwise), is inconsistent with the hypothesis of a recent *emm4* to *emm12* transmission event and argues for emergence of macrolide-resistant *emm4* and *emm12* lineages into Iceland not being directly related.

Another possibility is that the emergence and expansion of the macrolide resistant *emm* types contributing to the three epidemic waves that occurred in Iceland between 1998 and 2008 was driven by changes in antibiotic usage. Antimicrobial consumption for macrolides in Iceland was fairly constant from 1997 to 2009, with mean annual outpatient usage ranging from 1.85 defined daily doses per 1000 inhabitants per day in 1998 to 1.25 in 2009 (44). Over this period there was a gradual decrease in the use of short-acting macrolides (i.e. erythromycin), and a corresponding increase in the use of intermediate (i.e. clarithromycin) and long-acting (i.e. azithromycin) macrolides, but no year-to-year dramatic shifts occurred (Figure S2). The detected macrolide-resistant *S. pyogenes* isolates increased 12.8-fold between 1998 and 1999 (from 34 to 434 isolates, nearly all *emm4*) and increased 4.3-fold between 2007 and 2008 (from 65 to 281 isolates, nearly all *emm6*). The lack of any substantial change in macrolide usage corresponding with these dramatic increases in detected macrolide resistant isolates is inconsistent with emergence and expansion being driven by antibiotic selective pressure.

Although beta-lactam susceptibility testing was not done for all of the Iceland macrolide-resistant *emm12* isolates, it is likely that all 327 isolates that have the PBP2X Met593Thr amino acid substitution have reduced beta-lactam susceptibility with ~two-fold increased MICs for penicillin G and ampicillin. This idea is supported by the findings that there was two-fold increased penicillin G and ampicillin MICs for the five tested *emm12* isolates temporally spread over the first (1999) and second (2004) peaks of macrolide-resistant infections, and the isogenic PBP2X Met593Thr substitution in the *emm89* genetic background that demonstrated this single nonsynonymous A to C nucleotide change in *pbp2x*/single amino acid Met to Thr change in PBP2X is sufficient to increase penicillin G and ampicillin MICs two-fold. It needs to be made clear that none of the isolates tested had MICs meeting the *in vitro* definition for penicillin or ampicillin resistance (EUCAST Clinical Breakpoint Tables v10.0: benzylpenicillin resistant >0.25 μg/ml). It is noteworthy that the PBP2X Met593Thr substitution is (along with the PBP2X Pro601Leu (12)) only the second PBP2X amino acid change to be experimentally proven to reduce *S. pyogenes* beta-lactam susceptibility. A molecular understanding of how the PBP2X Met593Thr change alters beta-lactam susceptibility requires further investigation and would be aided by determination of a *S. pyogenes* PBP2X crystallographic structure.

One prevailing argument for why all bacteria have not evolved/acquired polymorphisms conferring resistance to any given antibiotic is that such resistance mutations result in organisms that are of reduced fitness in an environment that lacks that antibiotic (45–48). In such an environment, bacteria with fitness-reducing resistance mutations are, over time, out-competed by more fit, susceptible bacteria and consequently become less prevalent/go extinct in the population. To our knowledge, the identification in Iceland of the closely genetically related 327 *emm12* macrolide-resistant isolates is the largest population identified of *S. pyogenes* clinical isolates with a PBP2X substitution conferring reduced beta-lactam susceptibility that are clearly recent clonally-related descendants. Given that most of the isolates in this cohort come from pharyngitis patients, this indicates *S. pyogenes* strains with beta-lactam susceptibility-altering mutations in *pbp2x* can be readily transmitted and cause pharyngitis.

The identification of a large number of naturally-occurring GAS strains with *mef*(A)-*msr*(D) M phenotype macrolide resistance, in conjunction with a *pbp2x* nonsynonymous mutation producing a peptidoglycan synthesis transpeptidase (PBP2X) that confers reduced beta-lactam susceptibility, is concerning given that beta-lactams and macrolides are the first and second antibiotics of choice for treating *S. pyogenes* infections. Such strains are potentially stepping stones along the evolutionary path to true beta-lactam resistant GAS. The use of either beta-lactam or macrolide antibiotics could provide the selective environment that favors the survival of such strains, increasing the opportunity for the incremental accumulation of additional resistance-enhancing polymorphisms. This emphasizes the need for beta-lactam susceptibility monitoring of GAS and the need for a vaccine to prevent GAS infections.

## ACKNOWLEDGMENTS

This study was supported in part by the Fondren Foundation, Houston Methodist Hospital and Research Institute, and National Institutes of Health grants AI139369 and AI146771 (to J.M.M.).

---

**Figure S1.**
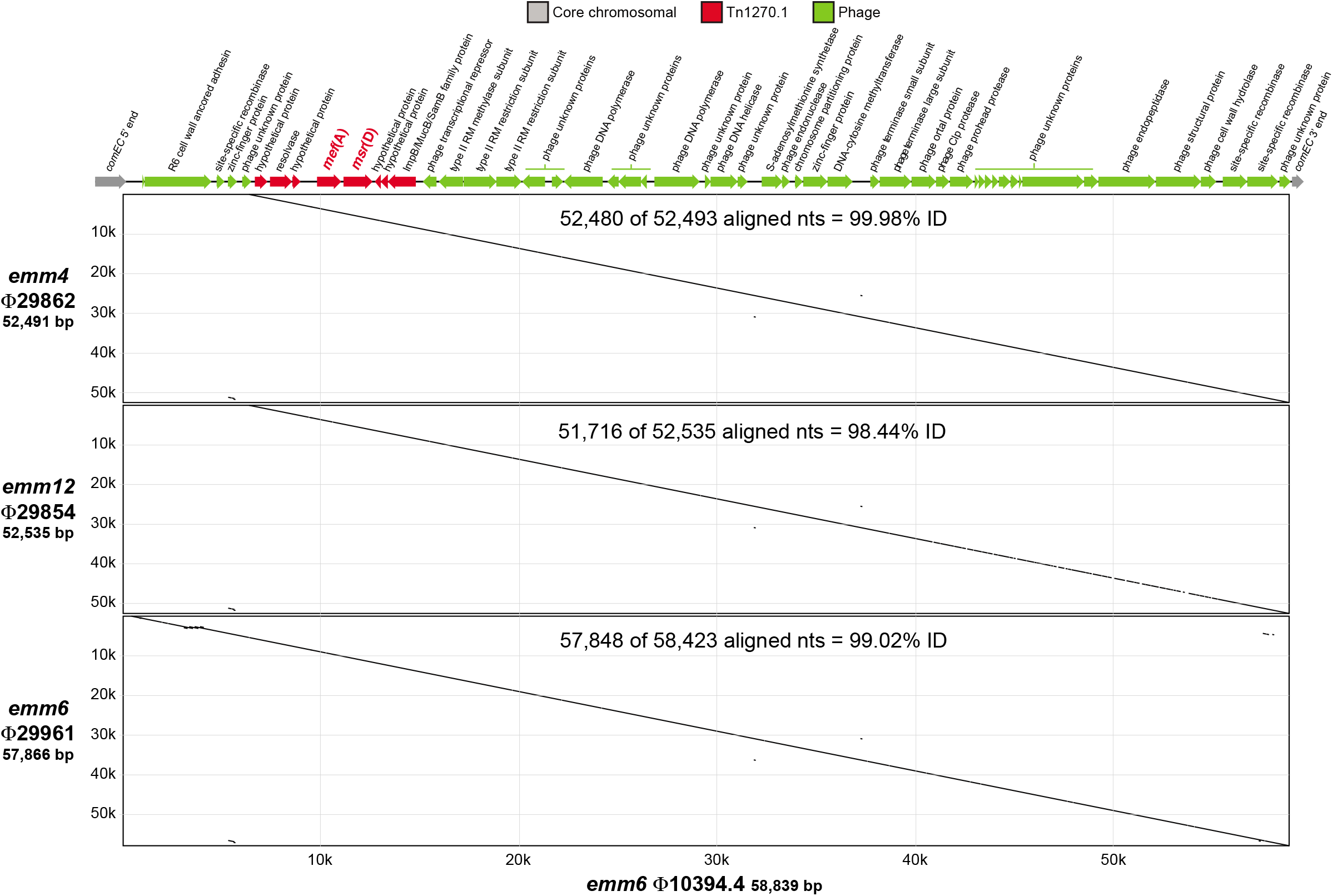
Composite mobile genetic elements (MGEs) encoding *mef*(A)-*msr*(D). Shown are dot matrix alignments of the *mef*(A)-*msr*(D) encoding composite MGEs identified in the Iceland erythromycin-resistant *emm* type 4, 6, and 12 clonally-related isolates with chimeric element Φ10394.4. Alignments were made with a 20 nt moving window at 95% identity. Core chromosomal, Tn1270.1, and phage gene content of the Φ10394.4 gene map are colored as indicated in the index.

**Figure S2.**
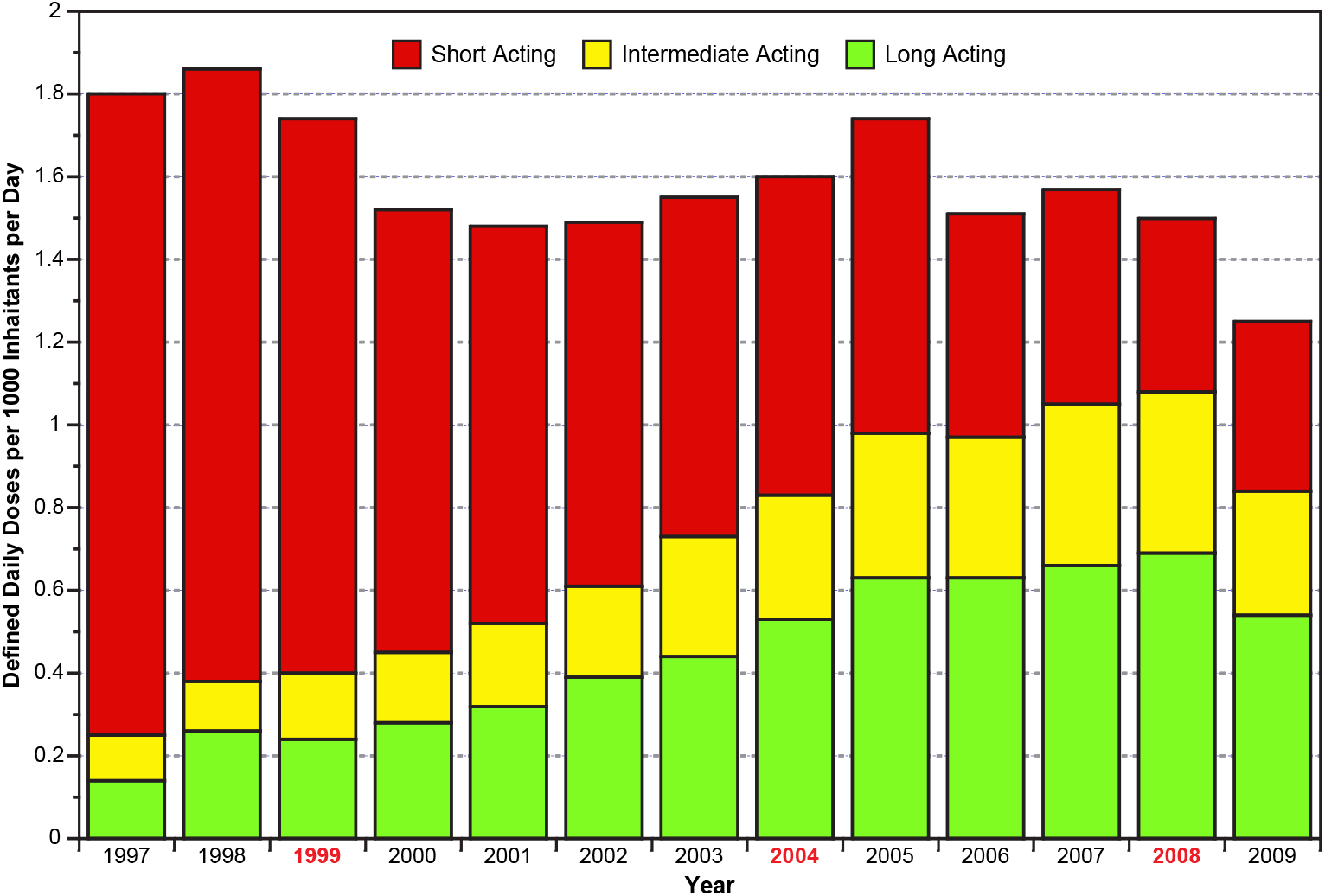
Annual outpatient consumption of macrolides in Iceland in defined daily dosages per 1000 inhabitants per day. The macrolides are categorized according to their plasma half-lives as short-acting (<4 hr, e.g. erythromycin), intermediate-acting (>4 but <24 hr, e.g. clarithromycin), or long-acting (>24 hr, e.g. azithromycin), as indicated in the index. The data was taken from the European Surveillance of Antimicrobial Consumption (ESAC): outpatient macrolide, lincosamide, and streptogramin (MLS) use in Europe (1997-2009), Table 2 and derived from the report (44).

